# Directed Network Discovery with Dynamic Network Modeling

**DOI:** 10.1101/074286

**Authors:** Stefano Anzellotti, Dorit Kliemann, Nir Jacoby, Rebecca Saxe

## Abstract

Cognitive tasks recruit multiple brain regions. Understanding how these regions influence each other (the network structure) is an important step to characterize the neural basis of cognitive processes. Often, limited evidence is available to restrict the range of hypotheses a priori, and techniques that sift efficiently through a large number of possible network structures are needed (network discovery). This article introduces a novel modeling technique for network discovery (Dynamic Network Modeling or DNM) that builds on ideas from Granger Causality and Dynamic Causal Modeling introducing three key changes: 1) regularization is exploited for efficient network discovery, 2) the magnitude and sign of each influence are tested with a random effects model across participants, and 3) variance explained in independent data is used as an absolute (rather than relative) measure of the quality of the network model. In this article, we outline the functioning of DNM and we report an example of its application to the investigation of influences between regions during emotion recognition. Across two experiments, DNM individuates a stable set of influences between face-selective regions during emotion recognition.

**New and Noteworthy:** In this article we introduce a new analysis method (Dynamic Network Mod- elling or DNM) which exploits *ℓ*_1_ regularization to perform efficient for network discovery. DNM provides information about the direction and sign (inhibitory vs excitatory) of influences between brain regions, and generates measures of variance explained in independent data to evaluate quality of fit. The method is applied to brain regions engaged in emotion recognition, individuating a similar network structure across two separate experiments.

## 1 Introduction

When we perform a task, such as recognizing a face, attributing mental states to others, or understanding a sentence, multiple brain regions are engaged (Ishai [2008], Gallagher and Frith [2003], Fedorenko and Thompson-Schill [2014]). Studying how these brain regions influence each other is an important step to understand the neural mechanisms underlying task performance. Influences between brain regions can be specific to the particular task a participant is performing. For example, face-selective brain regions might influence each other more when we recognize a face than when we recognize a scene. For this reason, we need a method that goes beyond measuring the presence of anatomical connections between regions, and to investigate the relations between the regions’ responses in the context of a specific experimental paradigm.

The direction of an influence can convey information about its function. For example, an influence from the ventral visual stream to prefrontal cortex is likely to convey bottom-up perceptual information to categorization and decision processes, while an influence from prefrontal cortex to the ventral visual stream is more likely to affect visual processing via top-down attentional selection(Buschman and Miller [2007]). Directed influences between brain regions can also contribute to characterize the functional role of a brain region by investigating how it receives inputs and conveys outputs to other regions with functional roles that are better understood.

Directed influences can be studied using temporal precedence: observing if earlier responses in a region contribute to predicting later responses in another region. Studying temporal precedence with functional magnetic resonance imaging (fMRI) presents unique advantages but also unique challenges. Data can be acquired noninvasively, with good resolution, and covering the entire brain, making fMRI well-suited to study long-range influences and investigate uniquely human aspects of cognition. At the same time, fMRI measures Blood-Oxygen Level Dependent (BOLD) signal, whose timing is affected by the local properties of vasculature. Adequate steps must be taken to control for the variability in BOLD timing between regions.

Depending on the evidence already available, different approaches to studying influences can be more or less suitable. In some cases, the previous evidence can be used to restrict a-priori the hypotheses about the influences between a set of brain regions, paving the way for confirmatory analyses. Often, however, limited evidence is available, and a very broad range of different influences are possible. Currently, the main techniques used to study influences between brain regions with fMRI are Granger Causality (GC, Roebroeck et al. [2005]) and Dynamic Causal Modelling (DCM, Friston et al. [2003]). Each of these techniques has important strengths, but also properties that may not be desirable in some data analysis contexts.

### 1.1 Granger Causality

In GC (Roebroeck et al. [2005]), the influence of one brain region on another is measured as a function of the variance in the responses in the latter region that is explained by earlier responses in the former, in addition to the variance explained by the latter region itself (see Appendix II for a more formal description). GC thus offers an intuitive measure of influences between regions which is not computationally costly to obtain. Nevertheless, GC has some disadvantages when applied to haemodynamic responses measured by fMRI. First, in its current form Granger causality is difficult to apply to the modelling of influences in different conditions within fast event-related designs. Separate models are used for the different conditions that need to be compared, and in fast event-related designs this would require breaking up the timeseries in short chunks that would impair autoregressive modeling. Second, since GC is based on variance explained, it cannot measure whether stronger responses in one region lead to stronger or weaker responses in another (i.e. the ‘sign’ of the influence), or even whether this feature of the influence is consistent across participants. In the end, GC may be prone to overfitting, especially when many brain regions are considered, because of the large number of parameters and the lack of techniques to penalize model complexity (regularization).

### 1.2 Dynamic Causal Modelling

In DCM (Friston et al. [2003]), the change in neural response in each brain region is modeled as a function of the stimulus and the input from other regions using ordinary differential equations (see Appendix III for a more formal description). Given a set of brain regions and a set of conditions, there is a fixed number of possible parameters. A candidate model is specified by providing, for each possible parameter, a 1 if that parameter will be included in the candidate model (whether a connection‘exists’ in that candidate model), and a 0 otherwise. The parameters that are included are then estimated with the expectation-maximization (EM) algorithm, and the best candidate model is chosen using Bayesian model comparison. The presence of condition-dependent influences makes DCM suitable for the analysis of fast event-related designs, and the use of priors mitigates overfitting acting as a form of regularization.

The proposed DNM approach adapts many of the strengths of DCM, but takes a different approach to two key challenges. The first challenge is network discovery. DCM is designed as a confirmatory technique, and therefore it is particularly suitable for choosing between models that have been identified on the basis of prior evidence. Searching through larger hypothesis spaces in DCM is computationally costly (but see Friston and Penny [2011] for an ingenious technique to increase speed); but more importantly, DCM computes the posterior probability of the best model, relative to the set of considered models. This estimate is most useful when researchers can be confident that their a priori hypothesis space includes most or all plausible models of the network. By contrast, DNM is designed as an exploratory or network-discovery approach. We propose that DNM can be used by researchers who cannot restrict their hypothesis space to a small set of models based on existing evidence, or who are unsure whether the limited temporal resolution of fMRI data is sufficient to reveal any robust inter-regional influences.

Whereas DCM compares full network models (each model is a description of the overall structure of the network), DNM assesses the variance in independent data explained by each individual parameter (or connection), using regularization to efficiently search all possible connections. A consequence of this difference is that whereas the best model chosen by DCM includes both the existing and the absent connections, DNM (like traditional null-hypothesis tests) makes claims about the existing connections, but not about the absent connections. On the other hand, DNM provides an absolute, not relative, estimate of the variance explained by each connection in independent data. This intuitive measure of the quality of the model is especially useful for exploratory or network-discovery analyses.

Existing packages in DCM employ free energy to assess quality of the model fitting to the data. Although this function is based in the same data used for model selection, it is not biased in favour of complex models.DCM also offers the option to calculate variance explained by each model, but using the same data used for model selection. As a consequence, the variance explained by the selected model is overestimated. By contrast, DNM uses variance explained in independent data. Similarly to free energy, this is not biased by model complexity. However, variance explained in independent data has an important additional asset: it is an accurate estimate of the absolute measure of goodness of fit. Variance explained in independent data can be used to how well the model fits the current data (for example, making clear when even the best model provides a relatively poor fit), but also to compare model fits across experiments and populations.Therefore we chose variance explained in independent data as a key measure to evaluate quality of fit in DNM.

The second challenge is how to combine evidence across participants who may have individual differences in the strength of each connection. In DCM, each candidate model specifies the existence and direction of connections, but not their value (Stephan et al. [2010]). That is, in DCM, a connection is deemed present if it explains variance, regardless of whether the parameters are similar (or even the same sign, that is ‘excitatory’ vs ‘inhibitory’) across participants. This approach grants DCM the flexibility to identify connections with highly variable strengths across participants, buthas consequences for the intuitive interpretation of the resulting model graphs. A high degree of variability in the parameter values across participants does not affect the probability of the model given the data. By contrast, DNM follows traditional null-hypothesis testing, in assessing the reliability of the magnitude (and sign) of each parameter across participants. This difference can be illustrated using methods for testing whether an experimental condition leads to a significantly greater response than baseline. In standard fMRI analyses, using a general linear model, a voxel is deemed to show a significant response if the magnitude (i.e. beta parameter) of response is reliable (similar in both magnitude and sign) across participants. A valid but different analysis would ask whether including the predictor for the experimental condition improves the fit of the model across participants (enough to compensate for the increased complexity of the model). In this case, the beta values might be highly variable, or even positive for some participants and negative for other participants, as long as they contribute enough to improving the model fit in each participant. Both of these analyses are valid, but address different questions. DCM takes an approach similar to the second analysis: when a model including is selected, we can infer that each included connection improved the quality of fit enough to compensate for the additional complexity, but the parameter value for that connection might be quite different across participants. This observation has been recently raised in the MEG literature using DCM (Pinotsis et al. [2016], Friston et al. [2016]), where it has been addressed using an approach based on hierarchical Bayesian models. In DNM, we adopt an alternative solution: using random effect statistical tests on the parameter values. This solution is intuitive and it interfaces well with the regularization approach to network discovery.

### 1.3 Dynamic Network Modelling

Given the considerations outlined in the previous sections, we set out to develop a new conservative method for exploratory analysis to model influences between brain regions in fMRI, that would meet a set of criteria: 1) control for the differences in the shape of haemodynamic responses in different regions, 2) provide a computationally efficient method for network discovery, 3) provide rigorous estimates of the magnitude and the sign of influences (whether they are inhibitory or excitatory), and 4) offer absolute measures of goodness of fit (rather than relative to other models) in independent data. To satisfy these desiderata, here we propose a novel approach to modeling dynamic influences between brain regions that builds on insights from GC and DCM and introduces some new ideas. We begin with deconvolution as in DCM, and proceed with a regularized vector-autoregressive modelling, obtaining a procedure that we refer to as Dynamic Network Modelling (DNM). This two-step procedure in which DCM-like deconvolution is followed by subsequent network modelling has been shown to successfully control for HRF shape (David et al. [2008]). Like GC, DNM models the effect on a region of the stimuli and of other regions with a system of linear equations. Like DCM, DNM includes parameters for fixed and condition-dependent connections as well as driving effects of the experimental conditions, and it can model independent connections in opposite directions between the same brain regions. Unlike both, DNM fits parameters with *ℓ*_1_ (Lasso) regularization, simultaneously addressing the overfitting problem and performing network discovery thanks to the sparsity encouraged by this regularization approach. Autoregressive parameters, condition-dependent effects and inter-regional influences are fit sequentially in a hierarchical model, yielding conservative estimates of the impact of the influences between regions. We assess the model using variance explained in left-out data to measure the quality of the model fit without a bias for complex models, and we use t-tests on the model parameters to identify influences that are consistent in sign and magnitude across participants.

### 1.4 Empirical Application

We adopted DNM to investigate the influences between brain regions involved in the recognition of emotional valence: an ideal case study because recent studies (Peelen et al. [2010], Skerry and Saxe [2014]) identified multiple brain regions encoding information about valence, and because recognition of emotional valence is likely to engage species-specific processes whose investigation necessitates noninvasive measurement. A brief glance at a person’s face provides rich information: about the person’s identity, age and gender, and also about their current emotional experience. Many brain regions are involved in extracting this information (Anzellotti and Caramazza [2015]). For example, the occipital and fusiform face areas (OFA and FFA) respond selectively to faces (Sergent et al. [1992], Kanwisher et al. [1997], Gauthier et al. [2000]) and encode information about many facial features, including the valence of the emotions communicated by facial expressions (Peelen et al. [2010], Furl et al. [2012], Skerry and Saxe [2014]). The posterior superior temporal sulcus (pSTS) and medial prefrontal cortex (MPFC) are sensitive to cues about another person’s emotional experience, whether conveyed in facial expressions or in body movements, vocal tones, or even abstract desriptions of events (Narumoto et al. [2001], Winston et al. [2005], Fusar-Poli et al. [2009], Peelen et al. [2010], Furl et al. [2012], Skerry and Saxe [2014]).

A key open question concerns the interaction between these two kinds of information (see Ishai [2008], Bressler and Menon [2010]): how does the facial form processing in OFA and FFA interact with the more invariant representations of emotional valence in pSTS and MPFC? One possibility is that recognition of emotion in facial expressions is accomplished mainly by successive bottomup feature extraction, building increasingly invariant representations at each stage. If so, one would expect influences mainly from OFA and FFA to pSTS and MPFC. Another possibility, however, is that higher-level representations of emotions influence the processing of the facial form. If so, one might expect that inter-regional influences include, or are even dominated by, top-down influences from pSTS and MPFC on the processing in OFA and FFA.

Structural connectivity analyses suggest that although OFA and FFA are connected, there are no direct (i.e. monosynaptic) tracts connecting pSTS to the OFA or FFA (Davies-Thompson and Andrews [2011], Ethofer et al. [2011, 2013], Gschwind et al. [2011], Pyles et al. [2013]). Direct anatomical connections between MPFC and OFA/FFA are also unlikely. Nevertheless, the magnitude of spontaneous activity during rest (i.e. in the absence of any emotional stimuli) is correlated between FFA and pSTS (Turk-Browne et al. [2010]) suggesting at least indirect connectivity between these regions.

We applied DNM to data from two independent experiments in which participants viewed brief (4 sec) movies depicting a stranger’s positive or negative emotional experiences. Multi-voxel pattern analyses of the first dataset suggested that information about the valence of these expressions is present in OFA, FFA, pSTS and MPFC (Skerry and Saxe [2014]). We therefore used these data, without a-priori restrictions on the connectivity profiles, to ask (1) how these regions interact during the recognition of emotions, and (2) whether DNM explains additional variance in independent data (as compared to a standard GLM) and produces reliable results (replicating connectivity profiles across different datasets).

## 2 Materials and Methods

### 2.1 Preprocessing and Deconvolution

Given a set of *n* ROIs, the mean BOLD signal is extracted from each of the ROIs, and detrended with the SPM function ‘spm detrend’ (http://www.fil.ion.ucl.ac.uk/spm/). Model-based deconvolution was performed as in David et al. (2008), using DCM software based on a modification of the balloon model to obtain estimates of the shape of the haemodynamic response function (HRF; Friston et al. [2003]). This deconvolution method was shown to control successfully for variability in the HRF shape between different brain regions in fMRI data, leading to fMRI-derived estimates of connectivity that correspond to those obtained with intracortical EEG (David et al. [2008]). For this reason, we do not provide additional validation tests here for this deconvolution approach.

### 2.2 Interaction Model

Deconvolution yields a deconvolved timeseries *z*_*i*_(*t*) for each ROI *i* = 1,…, *n*. The vector of deconvolved timeseries **z**(*t*) = [*z*_1_(*t*);…,*z*_*n*_(*t*)] is modelled with a bilinear vector autoregressive model. A bilinear model can model condition-dependent effects even in the context of fast event-related designs, and it does not require integration to estimate parameters thus reducing the computational costs. We have therefore

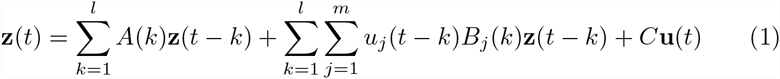

where *k* = 1…,*l* are the time lags between predicted and predictor responses, the matrices *A*(*k*) contain parameters for fixed influences at lag *k*, *u*_*j*_(*t*) is 1 if condition *j* is presented at time *t* and 0 otherwise, the matrices *B*_*j*_(*k*) contain parameters for condition-dependent influences in condition *j* at lag *k*, and *C* contains parameters for the effect of the conditions **u**(*t*).

### 2.3 Model Fitting

Two fundamental considerations were made in the context of model fitting. First, fitting all model parameters in a single stage would lead any shared variance between parameters for the influence of the regions on themselves (autoregressive parameters) and parameters for the influences between different regions to be split equally among them. Timecourses of fMRI data show a high degree of temporal autocorrelation (Woolrich et al. [2001]), and fitting all parameters in a single stage could attribute part of the variance explained by autocorrelation to influences between regions, leading to exceedingly liberal estimates of the influences between regions. For this reason, in a first stage the autoregressive parameters are fit, subsequently the effect of conditions, and in the end the parameters for the influences between different regions are estimated. Note that this approach is conceptually analogous to the practice in GC of assessing the reduction in residual variance introduced adding a region *A* as predictor to an autoregressive model modelling the response in a region *B* as a function of itself at an earlier time. This procedure is made explicit in the following set of equations:

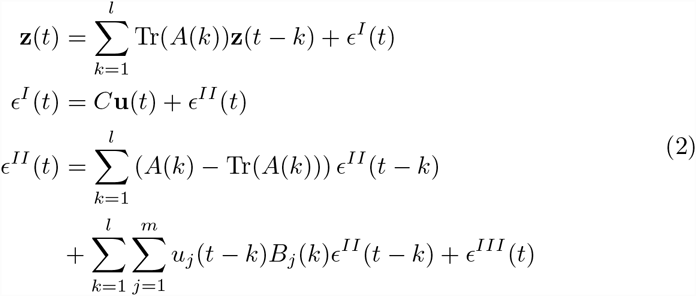

where 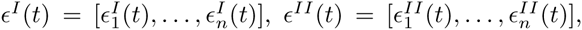 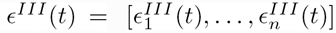are the residuals for the three modelling stages and the *n* ROIs. The first equation implements an autoregressive model, the second equation a linear model with conditions as predictors, and the final equation models the remaining variance as a function of the influences between different brain regions. The time lag *k* can be determined using the Akaike Information Criterion (AIC), but in practice given the high number of parameters the selected *k* is equal to 1. The advantage of fitting parameters hierarchically is that the estimates of the influences between regions are conservative. In this approach, autocorrelation within a timecourse is seen as a potential confound. If the main interest of a study instead is to estimate the extent of autocorrelation, fitting autoregressive parameters in a first separate stage could be too liberal. One potential cost of this hierarchical fitting procedure is that estimates of the shape of the HRF are generated before fitting the infulence parameters rather than simultaneously. However, previous studies found that the HRF could be estimated accurately even with such a sequential procedure (David et al. [2008]).

Second, the large number of parameters (especially when studying a large set of ROIs) calls for adequate techniques to reduce the problem of overfitting. For this purpose, a wide variety of regularization techniques are available in the literature (Hanke and Hansen [1993]). In keeping with the aim to provide conservative estimates of the influences between regions, the first stage of parameter fitting (concerning the autoregressive parameters) is not regularized, and parameters are estimated with OLS (Ordinary Least Squares). This enables the autoregressive component of the model to remove all the variance it can explain. Regularization is only employed for the last stage of parameter fitting, to reduce overfitting of the parameters describing the influences between brain regions. Among the many methods available for regularization, *ℓ*_1_ regularization has the amenable property of allowing for sparsity, and is achieved by minimizing the error function

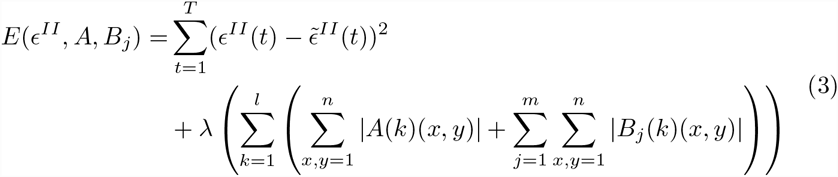

where *T* is the total length of the timeseries and

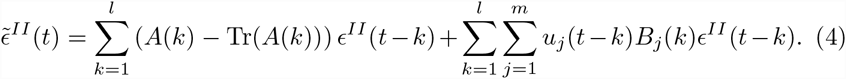

Note that the left part of equation [3] is the sum of square errors used in OLS, while the right part penalizes the use of parameters proportionally to *λ* times the sum of their absolute values. It is important to note that the error function in *ℓ*_1_ regularization is a sum of convex functions, and it is therefore convex (see Appendix IV). This implies that the error function has a single minimum, greatly facilitating parameter estimation. The value of *λ* is itself determined with a leave-one-out cross-validation procedure. Unlike *ℓ*_2_ regularization, *ℓ*_1_ regularization does not favor widely distributed and small parameter values, therefore it is also suitable to identify sparse networks in which only a subset of the influences are non-zero. Since *ℓ*_1_ regularization yields a convex error function and is suitable to identify sparse networks, it is an efficient tool for network discovery.

### 2.4 Experimental Materials and Procedures

Two different experiments testing emotion recognition from visual stimuli were completed with two different groups of participants. In each experiment, videos prompting the attribution of an emotion to a target were shown to participants while fMRI data were collected. The blood-oxygen level dependent (BOLD) timeseries measured were used to model connectivity between face selective regions and regions encoding information about emotional valence (OFA, FFA, pSTS and MPFC, see Skerry and Saxe [2014],Kliemann et al. submitted).

#### 2.4.1 Experiment 1: Stimuli and Task

A total of 26 volunteers took part in experiment 1 (all right handed, n = 10 female, age range: 19-44, mean = 26.25, SD = 6.12). All participants had normal or corrected to normal vision and no history of neurological or psychiatric disorders. Participants gave informed consent in written form in line with the requirements of MIT’s institutional review board.

The experiment consisted of 8 runs, a face localizer, and a theory of mind localizer (Dodell-Feder et al. [2011]; stimuli are available at http://saxelab.mit.edu/superloc.php). The theory of mind localizer was not analyzed for this article. In the 8 runs, participant viewed videos of faces expressing an emotion (‘expressions condition’) and of simple geometric characters experiencing an event eliciting emotions (‘situations condition’). Participants were asked to rate on a 1-4 scale after each video the intensity of the emotion experienced by the characters. Face stimuli involved a close-perspective view on single entity, therefore they were presented at 7.8 × 7.4° of visual angle, while the context
animations were presented at 16.7 × 12.5°. Videos of facial expressions were obtained from from movies, producing a set of relatively naturalistic stimuli, achieving a balance between external validity (see Zaki and Ochsner [2009], Spunt and Lieberman [2012]) and experimental control. Each trial consisted of the presentation of a 4s video followed by a 1.75s response screen and a 250ms blank. A fixation cross of variable duration (0-14s) was presented between trials.

In the face localizer, participants viewed videos of children’s faces and of moving objects (from Pitcher et al. [2011]). 30 videos were shown for each condition, grouped in blocks of 6. Participants had to perform a 1-back task detecting repetitions of identical videos. Each video lasted 3s, and was followed by a blank screen presented for 333ms, for a total block duration of 20s. A 2s blank was shown between every two blocks, and 12s blocks of fixation were included at the beginning, middle and end of the run. Participants were shown 2 localizer runs, each lasting 5 minutes. The order of conditions was counterbalanced within runs, across runs, and across participants.

#### 2.4.2 Experiment 2: Stimuli and Task

A total of 28 volunteers took part to experiment 2 (all right handed, 11 female, age range: 21-33, mean = 26.6, SD = 4.2). All participants had normal or corrected to normal vision and no history of neurological or psychiatric disorders. Participants gave informed consent in written form in line with the requirements of MIT’s institutional review board. The experiment was composed of 8 functional runs (lasting 372s each) and a face localizer (184s) (Hariri et al. [2000]).

In the experimental runs, the facial expression videos from Experiment 1 were presented. Unlike in Experiment 1, however, participants were asked to perform two different judgements on the facial expression videos. In half of the trials, they were asked to assess the valence of the emotion (positive or negative), in the remaining trials they were asked to assess the age of the person in the video (younger than 40, older than 40). Each trial consisted of the presentation of a screen indicating the task to be performed (1s), a blank (4-12s, mean = 8s), a video (4s), an additional blank (250ms) and a screen showing a plus and a minus arranged horizontally as a response cue (1.75s). Participants pressed the left or right button to select the plus (for positive emotions or for faces of people over 40) or the minus (for negative emotions or for faces of people under 40). At the end of each run, a 12s blank screen was presented. Participants were excluded if they performed below 83% accuracy in two or more runs, a criterion established before starting fMRI analysis. Data from three participants (1 female) were excluded due to this exclusion criterion.

The face localizer consisted in the presentation of 30s blocks of geometric shapes and blocks of faces. Each block consisted of 15 2s trials in which participant had to match the expression of a face (or the shape) in the upper part of the screen with one of two expressions (or shapes) in the lower part of the screen.

#### 2.4.3 Data acquisition

Data acquisition parameters were identical in the two experiments. Data were acquired with a 3T Siemens Tim Trio scanner in the Athinoula A. Martinos Imaging Center at the McGovern Institute for Brain Research at MIT, using a Siemens 32-channel phased array head coil. A high-resolution (1mm isotropic voxels) T-1 weighted MPRAGE anatomical scan was followed by functional scans using a gradient-echo EPI sequence sensitive to blood-oxygen-dependent (BOLD) contrast (repetition time [TR] = 2s, echo time [TE] = 30ms, flip angle = 90°, voxel size 3 × 3 × 3mm. 32 axial slices of 64×64 voxels aligned with the anterior/posterior commissure were acquired in each volume, covering the whole-brain except the cerebellum.

#### 2.4.4 ROI definition and Interaction Modelling

Data from the face localizers were preprocessed using SPM8 running on MATLAB R2010b and custom in-house software, and were modeled in both experiments with a standard GLM with regressors for each stimulus category. Regions of interest for the occipital face area (OFA), fusiform face area (FFA) and posterior superior temporal sulcus (pSTS) were defined in individual participants using group-defined search spaces (Figure 1). 1 of the 26 participants in Experiment 1 did not complete the face localizer, in one OFA and FFA could not be localized, and in 1 pSTS could not be localized. In these cases ROIs were derived from group-level activation maps obtained from the localizers of the other participants. In addition to these face-selective ROIs, a region of interest was defined for MPFC at the group level, using a 12mm radius sphere centered in the peak of accuracy for classification of the valence of emotions in the study by Skerry and Saxe Skerry and Saxe [2014], at MNI coordinates [-2, 50, 34].

**Figure 1:**
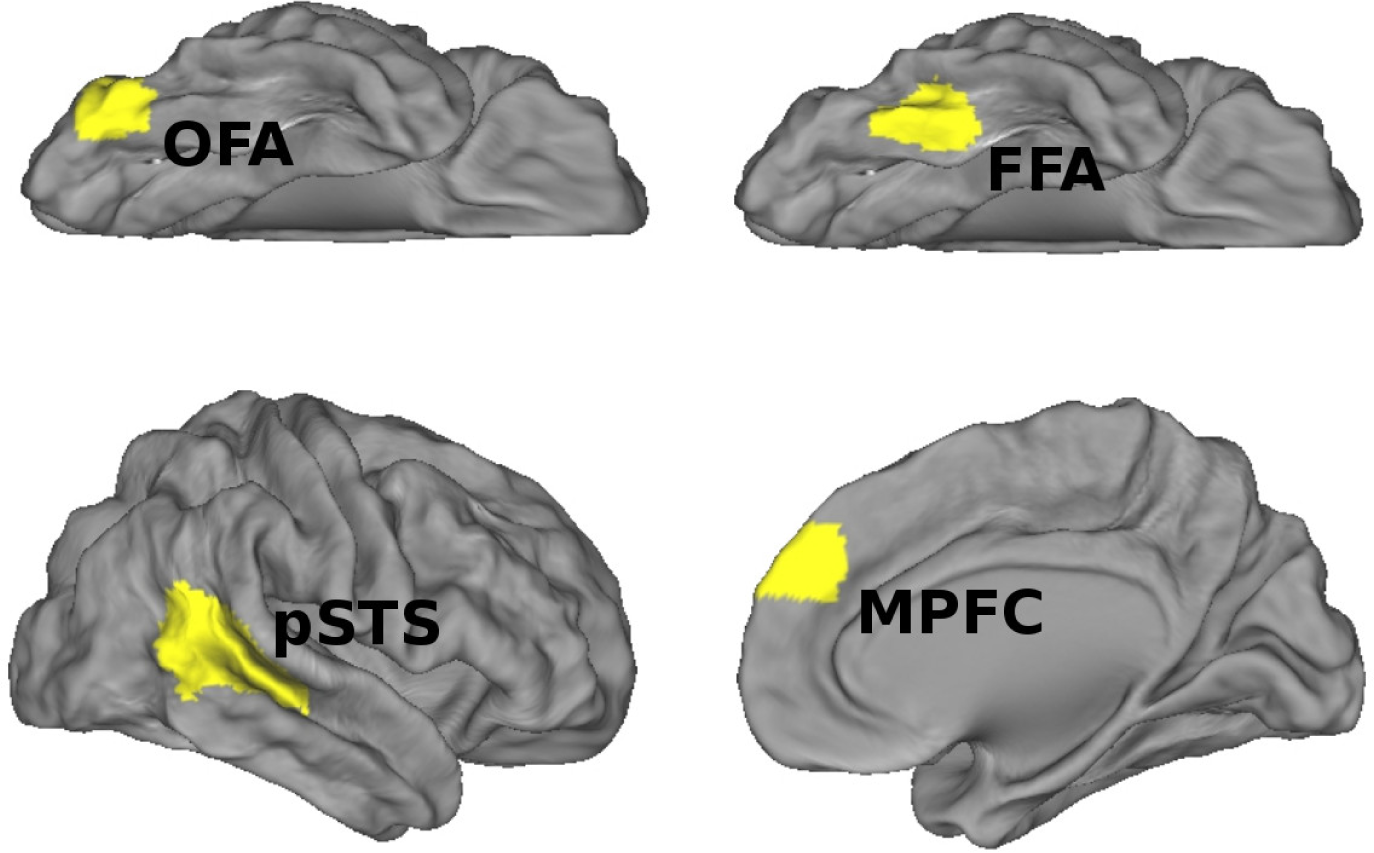
Group-level search spaces for the regions of interest.

Interactions between the ROIs’ mean timecourses were modelled using the method described in sections ‘Interaction Model’ and ‘Model Fitting’ above. In Experiment 1, faces and situations were used as conditions, in Experiment 2, age task and valence task were used as conditions.

## 3 Results

### 3.1 Experiment 1

The inter-regional influence parameters substantially improved the fit of the model, when added to the hierarchical model in the final stage. In the deconvolved timeseries, the influence parameters explained an additional 32.88% (SEM 2.02%) of the variance in left-out runs. A similar but weaker result was obtained without deconvolution, just predicting the original BOLD signal (additional variance 12.27%, SEM 0.9%, see Appendix I - Figure 3).

Network structure (Figure 2 A, B) was calculated modeling the deconvolved timeseries in the ROIs with a bilinear model and running a t-test on model parameters (see Materials and Methods for more details). Among face-selective regions, significant connections were found from the OFA to the FFA (*t*(25) = 2.76, *p* < 0.05), trends for influences were found from the FFA to the pSTS (*t*(25) = 2.00, *p* = 0.056) and from the pSTS to the OFA (*t*(25) = 2.05, *p* = 0.051). In addition, a significant influence from the OFA to the MPFC was observed (*t*(25) = 2.15, *p* < 0.05).

**Figure 2:**
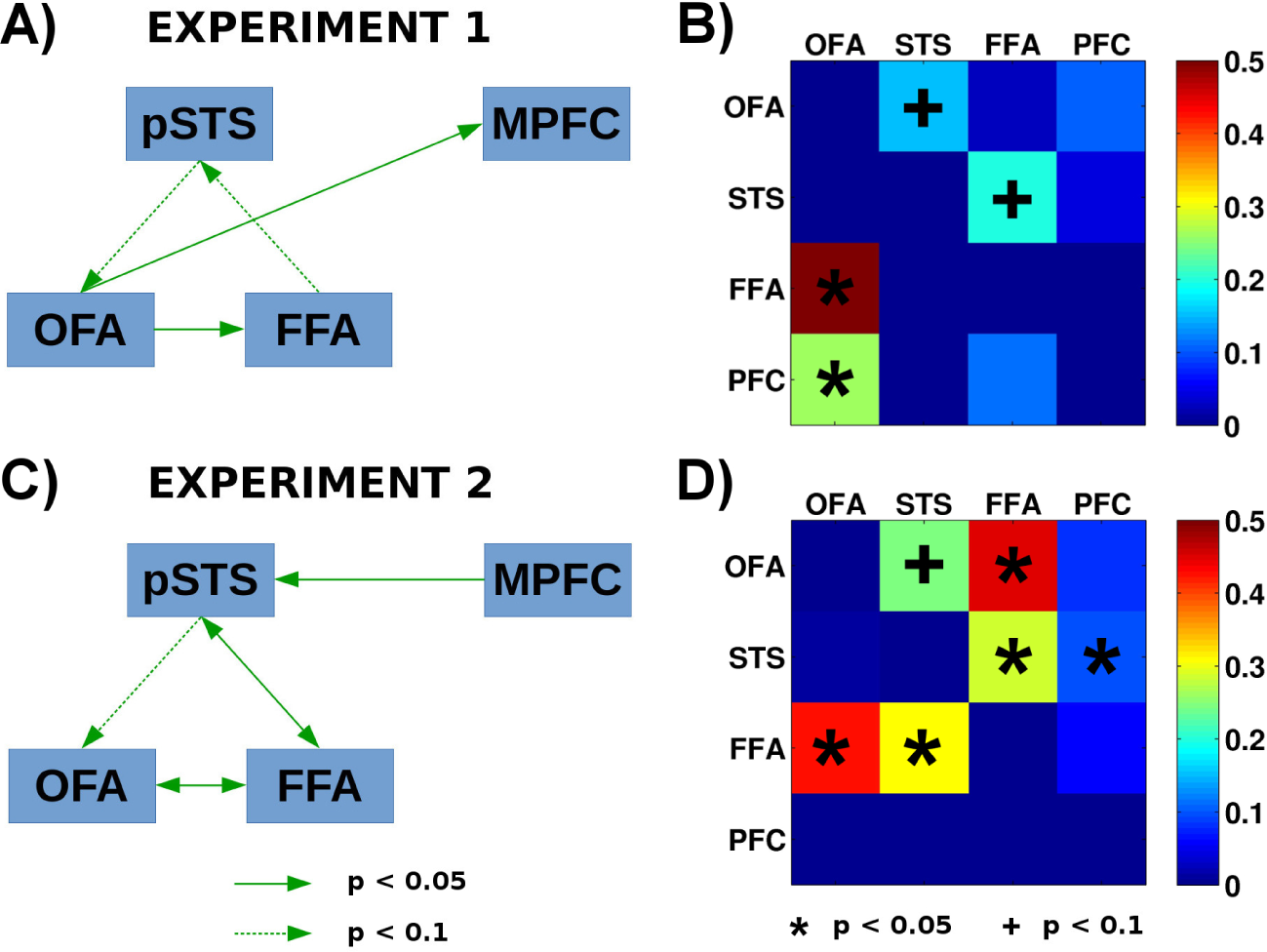
A) Connectivity diagram for experiment 1; B) color-coded matrix of parameter values for the influences between regions in experiment 1; C) Connectivity diagram for experiment 2; D) color-coded matrix of parameter values for the influences between regions in experiment 2.

**Figure 3:**
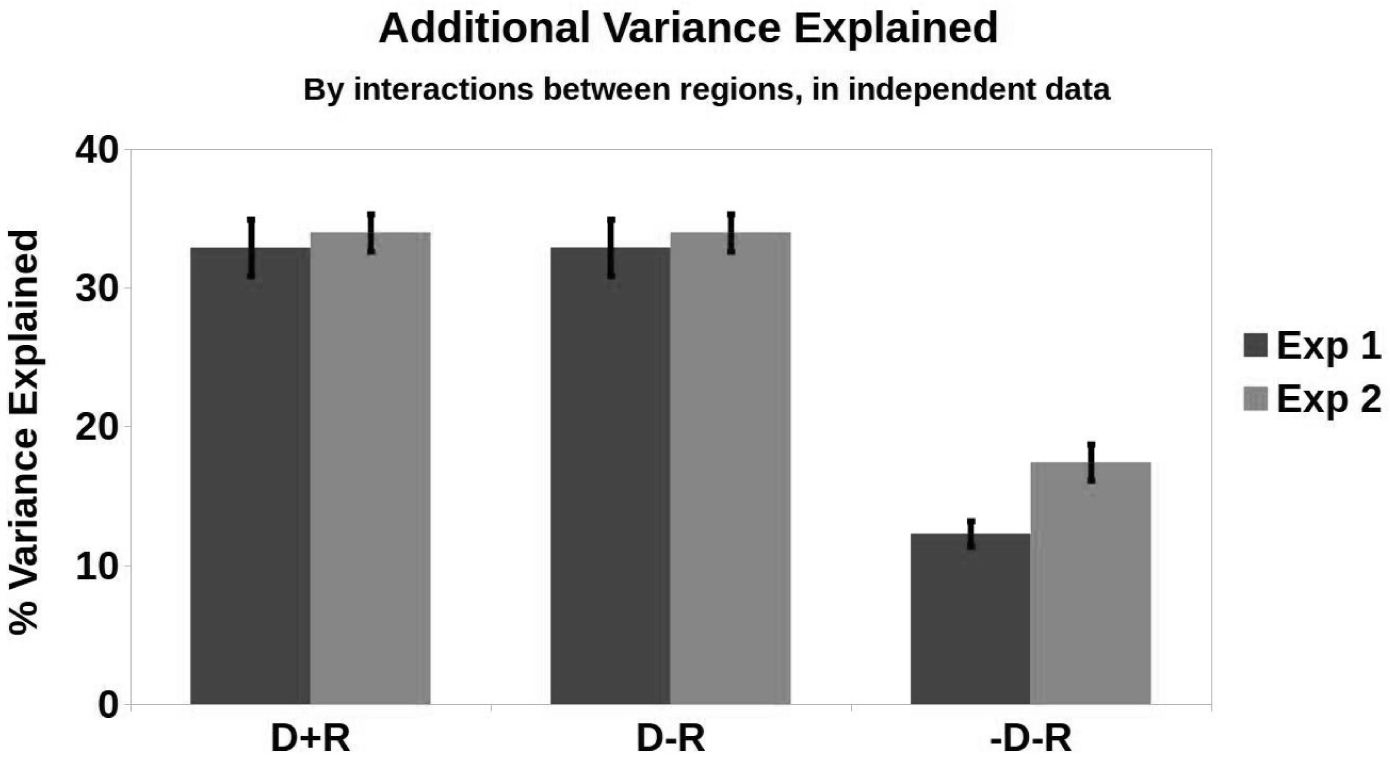
Additional variance explained in independent data by the influences between regions. The model using both deconvolution and regularization is compared to a model using deconvolution but not regularization and to a model using neither deconvolution nor regularization.

All condition-dependent influences were non-significant (all *p* > 0.1). Furthermore, a model without condition-dependent influences between regions was fit to the same data and produced the same results (Appendix I Figure 4). A model without regularization also generated similar results (Appendix I - Figure 5).

**Figure 4:**
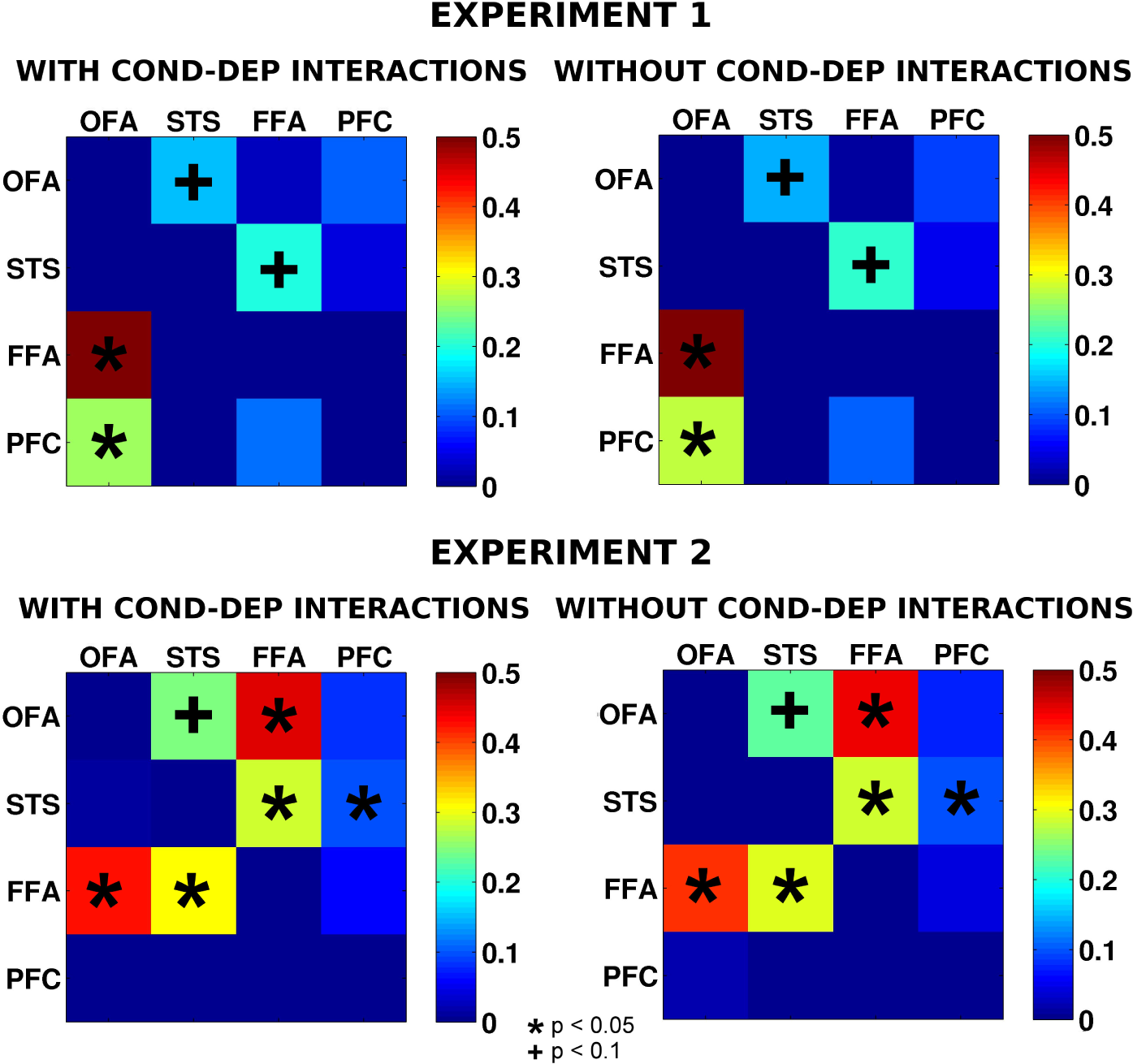
Comparison between parameter values for a model with and without condition-dependent influences between regions. Condition-dependent influences do not have a noticeable impact on the results with these data.

**Figure 5:**
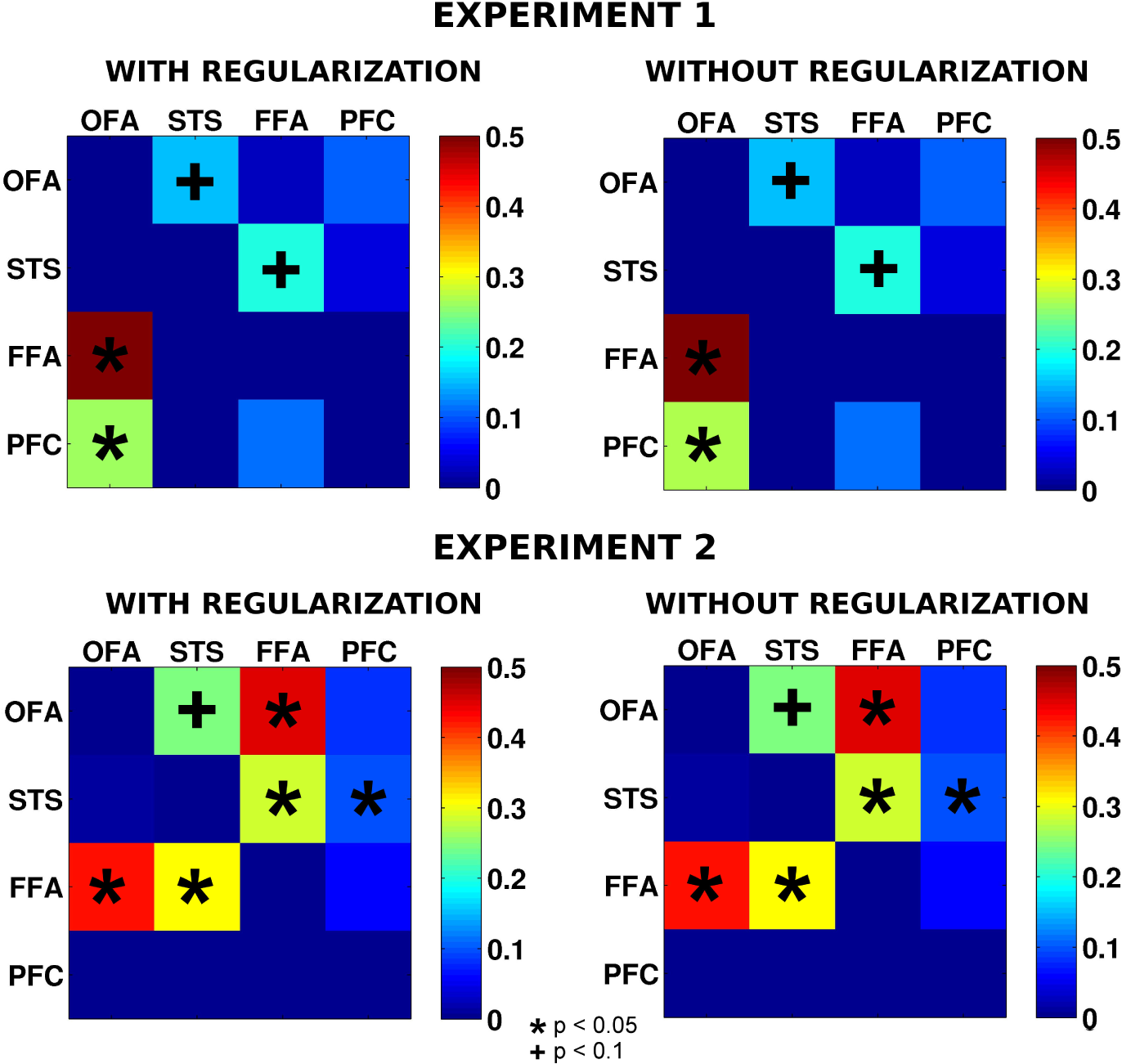
Comparison between parameter values for a model with and without regularization. Regularization does not have a noticeable impact on the results with these data.

On the other hand, deconvolution does appear to be critical. We compared DNM in deconvolved data to DNM in the original BOLD timeseries (Appendix I - Figure 6). In the original BOLD timeseries, multiple connections appear to be bidirectional (e.g. between OFA and FFA, pSTS and OFA, and PFC and pSTS), but after deconvolution (likely clarifying the relative timing of neural response in these regions), all of these connections appeared unidirectional.

**Figure 6:**
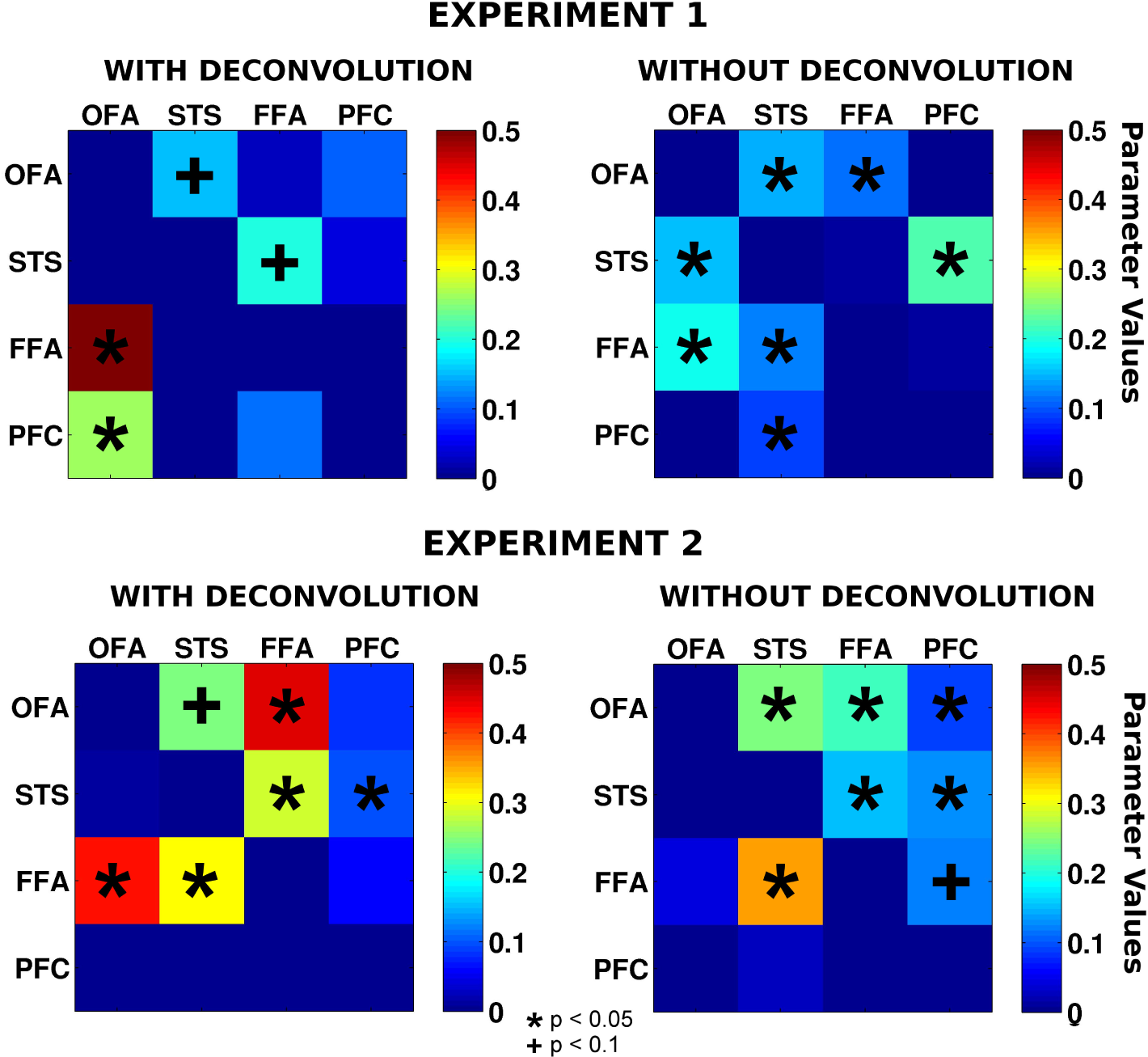
Comparison between parameter values for a model with and without deconvolution. Controlling for differences in haemodynamic responses between regions with deconvolution changes the resulting parameters.

### 3.2 Experiment 2

As in experiment 1, the inter-regional influence parameters (Figure 2 C, D) explained substantial additional variance in the deconvolved timeseries (33.97%, SEM 1.35%) and even in the original BOLD signal (17.4%, SEM 1.28%, see Appendix I - Figure 3). These values are remarkably similar to the results of experiment 1, suggesting that the importance of influence parameters is highly replicable, in this model, across participants and groups.

Network structure was calculated modeling the deconvolved timeseries in the ROIs with a bilinear model and running a t-test on model parameters (see Materials and Methods for more details). Among face-selective regions, in line with experiment 1, significant connections were found from the OFA to the FFA (*t*(24) = 4.50, *p* < 0.001), from the FFA to the pSTS (*t*(24) = 2.17, *p* < 0.05), and a trend was observed for the influence from pSTS to OFA (*t*(24) = 1.72, *p* = 0.098).

In addition, we observed influences from the FFA to the OFA (*t*(24) = 4.28, *p* < 0.001) and from the pSTS to the FFA (*t*(24) = 2.37, *p* < 0.05), as well as from the MPFC to the pSTS (*t*(24) = 2.69, *p* < 0.05). There also appeared to be a weak decrease in the influence from OFA to FFA, in both conditions (age condition: *t*(24) = −2.4, *p* < 0.05, emotion condition: *t*(24) = −1.9, *p* = 0.07), possibly related to shared variance between the condition-specific influence parameters and direct effects of the experimental stimulus modelled in an earlier stage of the hierarchical model. No other condition specific influences were observed.

A direct comparison of the results in the two experiments using unpaired samples t-tests did not yield any significant differences; a trend was observed for the influence from OFA to MPFC to be somewhat stronger in Experiment 1 (*t*(49) = 1.91, *p* = 0.0625).

## 4 Discussion

When participants watched naturalistic dynamic videos of emotional expressions, we observed reliable influences in cortical regional responses from OFA to FFA, FFA to pSTS, and pSTS to OFA. All three influences were positive (i.e. ’excitatory’) and replicated across two experiments, in different groups of subjects. In addition, the first study observed influences from OFA to MPFC, while the second study observed a reciprocal influence from FFA to OFA. Differences between studies are harder to interpret and discussed further below. More generally, we introduce a novel method for assessing dynamic influences between cortical regions that provides many key advantages over existing techniques in the context of network discovery.

We sought to measure directional, time-lagged influences between cortical regions, using functional MRI. An important challenge to using fMRI to study interations is the temporal resolution: the blood oxygenated dependent (BOLD) signal is inherently slow, and is typically measured at 0.5Hz. Other neuroimaging techniques, including MEG and EEG offer much higher temporal resolution than fMRI, and therefore the potential to study interregional influences at the true frequency of neural computation (e.g. >100 Hz). On the other hand, fMRI has substantially better spatial resolution. Poor source reconstruction is a serious challenge for studying interregional influences, because misattributing distinct signals to a common source could create a false appearance of an influence where none exists. Also, over the past decade, the cognitive functions of many cortical regions, localized with fMRI, have been intensively investigated, in terms of the magnitude and pattern of responses to stimulus categories. Further theory building will require understanding how influences between these particular cortical regions support transformation of information (i.e. processing).

Prior research investigating directional influences between brain regions in fMRI data has used two techniques: Granger Causality (GC, Roebroeck et al. [2005]) and Dynamic Causal Modelling (DCM, Friston et al. [2003]). While both techniques are powerful, and have been validated in some contexts (David et al. [2008]), both also have key limitations, especially in the context of network discovery, as identified in the introduction. Here we developed a new approach, building on the strengths of both GC and DCM, but also designed to satisfy novel criteria. Like DCM, we used deconvolution to address potential differences in the haemodynamic responses across regions, and simultaneously estimated parameters for interregional influences, and condition-dependent influences. These approaches address some limitations of the most common implementation of GC. On the other hand, compared to DCM, GC is conceptually intuitive and computationally efficient. In our case, we used *ℓ*_1_ regularized regression, which mitigates the overfitting problem and simultaneously offers an efficient approach to network discovery. Also, like GC, we aimed to provide a direct measure of variance explained, rather than the relative likelihood of a model (compared only to other models tested) provided by DCM.

Our approach (DNM) also goes beyond existing tools in multiple ways. First, we measure the variance explained by inter-regional influences, in independent left-out data. As an additional innovation, we used a step-wise regression, first greedily accounting for the stimulus-evoked responses and autocorrelations in the timeseries within region, before testing variance explained by inter-regional influences. In two experiments, we found that adding parameters for interregional influences to the model explained more than 30% of the remaining variance in the deconvolved timeseries of left-out runs. Evaluating variance explained in independent data is critical because it ensures that models are not overfitting the training data (variance explained in independent data is not biased in favor of more complex models), and also provides an absolute (not relative) measure of the goodness of fit of these models.

Second, we conduct statistical tests directly on the model parameters, across subjects, to preserve information about the sign of interregional influences (whether they are excitatory or inhibitory). In our data, all of the reliable interregional influences were positive (greater response in the source region predicted greater response, after a time lag, in the target region), but this method could also be used to detect and distinguish ’inhibitory’ influences (greater response in the source region predicts subsequently reduced responses in the target region). Furthermore, the method can be integrated with recent techniques for multivariate connectivity(Anzellotti et al. [2016])

The strongest inter-regional influence we observed was from OFA to FFA, in both experiments. Also, in both experiments, the evidence for this connection was stronger in deconvolved data, confirming that without deconvolution, hemodynamic differences may obscure the true direction of inter-regional influences. A connection from OFA to FFA is strongly predicted by the standard view of a a hierarchy of face processing from posterior to anterior regions in ventral temporal cortex (Lerner et al. [2001], Freiwald and Tsao [2010], Anzellotti et al. [2013]). In both experiments, influences from FFA to pSTS were found; studies measuring structural connectivity typically find no direct connection from FFA to pSTS, so the influence we observe here might be mediated by a third region not measured in this experiment (Davies-Thompson and Andrews [2011], Ethofer et al. [2011, 2013], Gschwind et al. [2011], Pyles et al. [2013]).

Both experiments also revealed an influence from pSTS to OFA (and to FFA in the second experiment). Prior evidence suggests that the pSTS contains a representation of facial expressions, including both dynamic facial movements generally, and emotional expressions in particular. The pSTS also contains a multimodal representation of emotions and facial movements. We hypothesize that the higher-level representations in pSTS, which are behaviourally relevant, modulate the processing of facial form in the OFA. An alternative account, in terms of predictive coding, would be that the representation of emotion in pSTS is used to predict of the perceived facial form in the ventral stream (see de Wit et al. [2010], Koster-Hale and Saxe [2013] for reviews). In a previous study, Furl et al. (Furl et al. [2014]) reported OFA-driven modulation of connectivity from V5 to STS. Our study extends these results by showing the presence of influences between motion processing and form processing involving both the OFA and FFA.

The measured influences with MPFC were relatively weak, and inconsistent across experiments. One caveat is that unlike the other three regions we tested, the MPFC was identified by a group ROI, rather than an individually-tailored ROI, so our measures of that regions’ timecourse may be less sensitive. On the other hand, it is possible that the inter-regional influences were actually different across the two experiments. In the first experiment, participants performed the same task on two different stimuli (faces, and abstract animations), and we observed an influence of OFA (possibly involved in face detection) on MPFC (involved in abstract emotion representation). By contrast, in the second experiment, participants performed two different tasks (emotion or age discrimination) on the same stimuli (faces), and we observed an influence of MPFC on pSTS, potentially modulating the relevance of emotional features. This possibility could be tested including both paradigms within an experiment with sufficient statistical power.

In general, modulatory connections might be particularly amenable to detection by fMRI, since they could have relative slow temporal frequency, compared with bottom-up information processing. The temporal resolution of fMRI is low, therefore the physiological bases of the signal modelled in this study are probably not monosynaptic connections between neurons. Temporal variation in the BOLD signal is correlated with local field potentials, and more specifically with the power of the gamma band frequency oscillations and with slow cortical potentials (*<*4Hz) (He et al. [2008]). The power of gamma band frequency oscillations and slow cortical potentials account for different parts of the BOLD’s variance (Scheeringa et al. [2011]). Slow cortical potentials are mostly driven by synaptic activity at apical dentrites in superficial layers of cortex (Goldring [1974], Mitzdorf [1985], Birbaumer et al. [1990], He and Raichle [2009]), and activity in superficial layers depends on long-range connections (He and Raichle [2009]). High power in the gamma band usually co-occurs with gamma synchronization within a region (Fries [2009]). It reflects engagement of the region in a task (Pulvermϋller et al. [1995]) and may need to be triggered by ‘modulatory network activation’, for instance top-down control (Fries et al. [2001, 2008], Bichot et al. [2005], Taylor et al. [2005], Womelsdorf et al. [2006]). Our results might depend on a combination of slow cortical potentials and gamma band power that reflect respectively low-frequency fluctuations in the activation state of a network and relatively faster sequences of responses due to engagement in a task.

In conclusion, we introduced an exploratory method (DNM) to model the dynamic, directed and signed influences between brain regions while controlling for inter-regional differences in the hemodynamic responses; parameter estimation with *ℓ*_1_ regularization addressed simultaneously the overfitting problem and the need for efficient network discovery. We applied DNM to the study of the influences between brain regions encoding information about emotional valence during emotion recognition, showing how DNM has key assets as compared to other available methods and gaining new insights on the influences between brain regions processing emotional valence.

## Acknowledgments

This study was supported by NIH Grant 1R01 MH096914-01A1 to Prof. Rebecca Saxe. Stefano Anzellotti was supported by a Postdoctoral Fellowship from the Simons Center for the Social Brain, Dorit Kliemann was supported by the Humboldt Foundation (Feodor Lynen Fellowship). We would like to thank the Athinoula A. Martinos Imaging Center at the McGovern Institute for Brain Research, MIT for providing scanning resources, and Rik Henson, Gabriele Anzellotti and Dimitris Pinotsis for comments on an earlier version of the manuscript.

## APPENDIX II Granger Causality

Let *a*(*t*), *b*(*t*) with *t* ∈ 1,…,*T* be two timeseries, for example the timecourses of Blood-Oxygen Level Dependent (BOLD) signal in two brain regions *A*, *B*. We can consider the autoregressive model

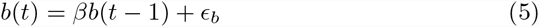

and the model

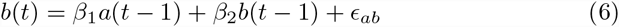

where *ϵ*_*b*_ and *ϵ*_*ab*_ are the residuals of the models. Here we focus for simplicity on a model between two regions that goes only 1 timestep in the past, but multiple regions and multiple timesteps in the past can also be considered. In Granger Causality, the influence from a brain region *A* to a brain region *B* is given by:

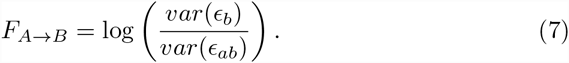

If earlier responses in region *A* explain additional variance in the responses of region *B*, the ratio *var*(*ϵ*_*b*_)/*var*(*ϵ*_*ab*_) is greater than one; the logarithm remaps the range of possible values from [1, +∞) to [0, +∞). The suggested approach to control for differences in haemodynamic responses between regions consists in calculating the subtraction:

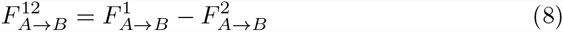

where 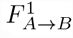 is the influence from *A* to *B* during condition *C*1, and 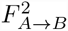 is the influence from *A* to *B* during condition *C*2.

## Appendix III Dynamic Causal Modelling

Let **z** = [*z*_1_(*t*),…,*z*_*n*_(*t*)] be a vector of neural responses at time *t*, where *z*_*i*_(*t*) is the response at time *t* in region *i* for each of *n* regions. In DCM, the change in neural responses ż (*t*) is modeled as

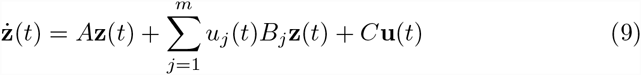

where *A* is a matrix of fixed connectivity parameters, *B*_*j*_ is a matrix of connectivity parameters during condition *j* for each of the *m* conditions, and *u*_*j*_(*t*) is 1 if condition *j* is presented at time *t* and 0 otherwise. The matrix *C* contains parameters for the effect of the conditions **u**(*t*) = [*u*_1_(*t*),…, *u*_*m*_(*t*)]. A candidate model *M* is specified providing matrices of binary values *Ã*, 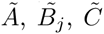 of the same size as *A*, *B*_j_, and *C*, so that in *M*, parameter *A*(*h, k*) is allowed to vary if and only if *Ã*(*h, k*) = 1. The parameters of candidate models are estimated with the expectation-maximization (EM) algorithm, and the best model is chosen using Bayesian model comparison (Friston et al. [2003]).

## Appendix IV Convexity of the error function in *ℓ*_1_ regularization

Let us consider 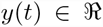 and 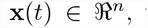 with *t* = 1,…, *T*. Let 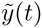 generated with a linear model 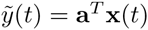 using parameters *a*_*i*_ = **a**(*i*), with *i* = 1,…, *n*. The error function given by *ℓ*_1_ regularization is

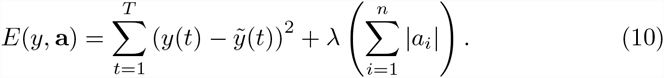

We can rewrite the error function as

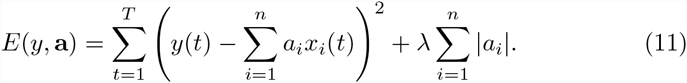

The function

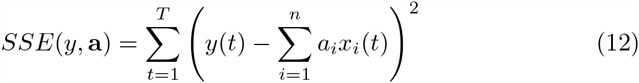

is convex in **a**. We also know that if two functions 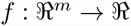 and 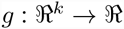 are convex, their sum 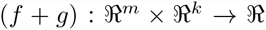 is convex. In fact, considering 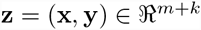 we have that

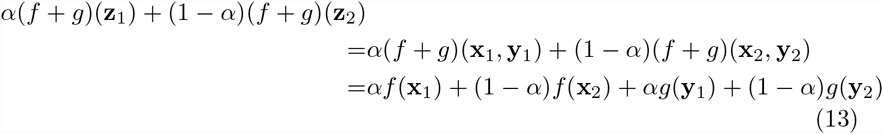

and since *f* and *g* are convex

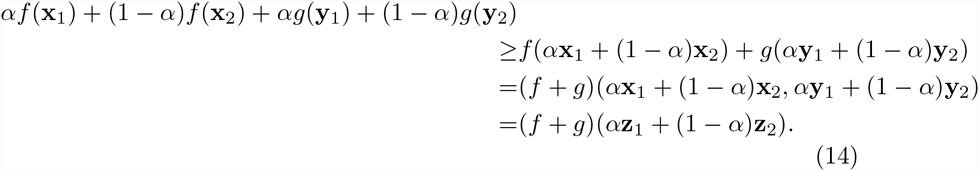

It follows that since |*a*_*i*_| is convex ꓯ*i* λ ≥ 0, also 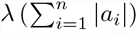 is convex. It can be shown with an analogous proof that given two convex functions 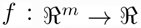 and 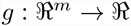 on the same domain, the function 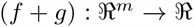 is also convex. Since the error function 10 is the sum of two convex functions on the same domain, it follows that it is itself convex.

Furthermore, the error function 10 is strictly convex, because it is a sum of a strictly convex function (the paraboloid given by the sum of square errors) and of a convex function (the penalization term). As a consequence, it has a unique minimum.

